# Structural semantic evolutionary distance (SSED) unifies the selection of cancer driver genes across macroevolution and tumorigenesis

**DOI:** 10.64898/2025.12.17.694808

**Authors:** Hongchen Ji, Qiong Zhang, Jianwei Wu, Fei Tian, Xiangxu Wang, Xiao Chai, Jiawei Chang, Mengdi Zheng, Xiao Li, Hong-Mei Zhang

## Abstract

The non-random, site-specific enrichment of somatic mutations in cancer driver genes (CDGs) suggests their emergence is governed by underlying evolutionary constraints. However, quantitative methods to define these constraints and their underlying principles remain underexplored. To address this, we introduce the Structural Semantic Evolutionary Distance (SSED), a metric leveraging the pretrained ESM-3 protein language model to quantify evolutionary divergence within a unified structural semantic space. Our analysis demonstrates that CDGs are subject to persistent structural semantic constraints across species, tolerating a significantly narrower range of structural semantic changes during evolution compared to non-CDGs. Crucially, clinically observed oncogenic mutations follow this same principle, favoring minimal structural perturbation as shaped by long-term gene evolution. Such mutations maintain core protein function while conferring a capacity for immune evasion, thereby driving clonal expansion. Guided by this “evolutionary constraint” framework, we successfully predicted and experimentally validated a previously uncharacterized oncogenic mutation, KRAS R135L, in bronchial epithelial cells. Furthermore, clinical cohort analysis demonstrated that SSED acts as an independent predictor of response to immune checkpoint blockade, offering information orthogonal to tumor mutational burden (TMB). This study unifies the evolutionary principles governing CDGs across macroevolutionary and microevolutionary (tumorigenesis) timescales, elucidates the balance between structural adaptability, functional conservation, and immune pressure, and identifies a novel predictive biomarker for cancer immunotherapy.

## Introduction

The development of cancers is generally considered to result from the interplay among genetic mutations, natural selection, and diverse selective pressures^1,2^. Among these factors, genetic alterations are regarded as the initiating events of tumorigenesis. Cancer driver genes (CDGs), including oncogenes (OGs) and tumor suppressor genes (TSGs), promote cancer development through gain-of-function or loss-of-function changes that endow cells with malignant phenotypes such as uncontrolled proliferation, resistance to apoptosis, and immune evasion^2^. From an evolutionary perspective, fragile genes or loci with high oncogenic susceptibility would be expected to be counterbalanced or eliminated by purifying selection. However, several CDGs (e.g. SMARCA4^3^, TP53^4^ and VHL^5^) remain highly conserved across multicellular organisms, suggesting that the presence of CDGs is maintained by strong evolutionary constraints and is therefore evolutionarily justified despite their oncogenic potential.

It is well established that many CDGs, such as SMARCA4 and VHL, participate in essential biological processes including cell-cycle regulation, development, and signal transduction^3,5^. These indispensable functions impose strict evolutionary constraints to maintain their structural and functional integrity. Some studies have proposed that oncogenic mutations may induce a “degenerative” phenotype, partially restoring behaviours reminiscent of unicellular ancestors^6^. However, this hypothesis does not adequately explain the complex and dynamic interplay between cancers and host metabolism and immune responses^7^. Notably, oncogenic mutations in CDGs often exhibit striking clustering at specific amino-acid residues and follow characteristic mutation patterns, such as KRAS (G12, G13, Q61)^8^, TP53 (R153H, R273H)^9^, and EGFR (L858R)^10^. The persistence of such clustering across genes and sites suggests that oncogenic mutations are not random events but are instead constrained by underlying evolutionary principles. This observation raises a central question: do oncogenic mutations merely follow the evolutionary rules governing CDGs, or is there a more general evolutionary logic that shapes the selection of oncogenic mutations?

Conventional analyses of gene conservation primarily focus on sequence similarity and substitution patterns, using metrics like dN/dS ratios and conserved site identification to assess selective pressures. These methods offer valuable insights into sequence-level evolutionary trends^11^. However, the structure and function of proteins are the ultimate substrates of selective pressure, serving as the critical link between genetic variation and organismal adaptation^12^. Advancements in protein language models have significantly enhanced our ability to characterize protein structure and function. For example, ESM3 can learn complex residue-level dependencies from amino acid sequences, predict protein structures and functions, and even generate novel proteins with defined functions^13^. These capabilities underscore its potential to capture deep structural semantics from amino acid sequences, thus providing a novel approach to studying protein evolutionary conservation at both structural and functional levels.

In this study, we introduce a novel structure-aware metric for quantifying evolutionary distance, termed Structure Semantic Evolutionary Distance (SSED). By leveraging structural semantic features generated from pretrained protein language models, SSED enables the assessment of conservation differences across species at the structural level. Using this approach, we demonstrate that CDGs are subject to stronger structural semantic constraints during evolution compared to non-CDGs. This “evolutionary constraint” also significantly influences the emergence and selection patterns of oncogenic mutations. Our findings show that oncogenic mutations are not entirely stochastic or deleterious events; rather, they occur within a constrained structural semantic space, extending the long-established evolutionary trajectory of CDGs and reflecting an intrinsic evolutionary driving force. Furthermore, based on cellular experiments and immunotherapy data, we show that this “evolutionary constraint” mechanism not only drives proliferative and immune adaptation but also critically influences responses to immunotherapy.

## Result

### 1 Structure Semantic Evolutionary Distance (SSED): A Structure-aware metric for protein evolutionary distance

To develop a protein evolutionary distance metric that integrates both sequence and structural information, we introduced the SSED method. This approach leverages per-residue feature vectors that encode structural and functional information, derived from the pretrained ESM3 protein language model. It quantifies structural semantic differences between sequences by computing the Minkowski distance (p=2) between corresponding feature vectors at each position of a multiple sequence alignment (Figure 1A).

**Figure 1.**
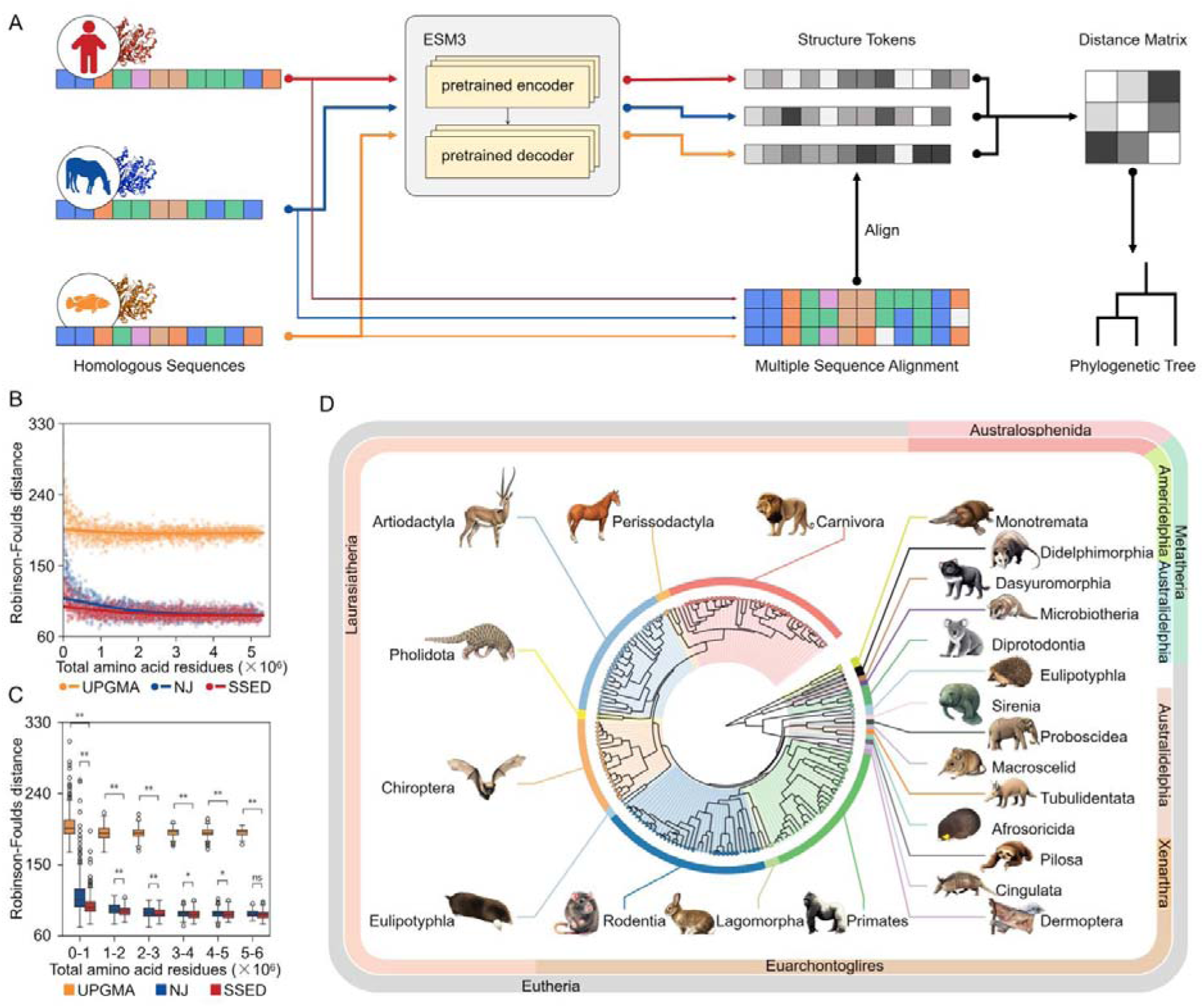
Workflow of the SSED method and evaluation of its performance in phylogenetic reconstruction. A: Flowchart of SSED calculation and phylogenetic tree construction. Homologous protein sequences were analyzed with the ESM-3 model to extract structural semantic representations (structure tokens). After multiple sequence alignment, pairwise SSED distances were computed and used to construct the phylogenetic tree. B: Comparison of phylogenetic reconstruction accuracy for the SSED method, Neighbor-Joining (NJ), and UPGMA methods. C: Quadratic fit of unweighted Robinson-Foulds distances for the three methods across different total amino acid sequence lengths. D: Comparison of mean unweighted RF distances for the three methods within segmented intervals of total amino acid sequence length (Mann-Whitney U test). E: Representative mammalian phylogenetic tree constructed using the neighbor-joining method based on the SSED distance matrix. ns: No significant difference, *: P<0.05, **: P<0.005.

To evaluate the effectiveness of SSED in inferring phylogenetic relationships, we conducted 3,000 simulations by randomly selecting sets of 10 to 3,000 protein-coding genes, with total amino acid lengths ranging from 15,652 to 5,315,823. Benchmarking against the reference phylogenies from TimeTree.org, we compared the phylogenetic trees reconstructed by SSED with those generated by classical distance-based methods, including the unweighted pair-group method with arithmetic means (UPGMA) and neighbor-joining (NJ). The results demonstrated that phylogenetic trees reconstructed with SSED exhibited significantly higher topological accuracy, as measured by the unweighted Robinson-Foulds distance, than those generated by UPGMA or NJ, irrespective of the total amino acid length (Figures 1B, C).

Using 16,959 protein-coding genes for alignment (Supplementary table 1), we reconstructed a phylogenetic tree of typical mammals based on SSED distances (Figure 1D), which showed high consistency with current established reference phylogenies, thereby validating the effectiveness of the SSED method. A notable discrepancy between the SSED-based tree and the established reference phylogenies concerned the classification of the order Eulipotyphla. This taxonomic group has been a long-standing challenge in mammalian phylogenetics. The SSED-based tree placed the families Soricidae and Erinaceidae as sister groups within the Afrotheria clade, whereas Talpidae was positioned at the basal node of the Laurasiatheria clade (Supplementary Figure 1). This topological arrangement aligns closely with early morphological studies, implying that the SSED method possesses sensitivity to protein structure and functional evolution, capturing stronger structural and functional similarities in cases of convergent evolution.

We further extended the SSED method to reconstruct phylogenetic relationships at the phylum level (Thermoproteota) and family level (Rhodobacteraceae). The results confirmed that SSED-based trees were more consistent with reference trees compared to those constructed using UPGMA and NJ methods (Supplementary Figure 2). Notably, the advantage of SSED became more pronounced when the total amino acid sequence length was shorter, suggesting that SSED-based phylogeny offers more accurate and robust evolutionary distance estimates under conditions of limited sequence information. This makes the SSED method particularly well-suited for analyzing evolutionary distances and relationships at the level of individual homologous proteins.

### 2 Widespread and persistent evolutionary constraints in CDGs

Using the SSED metric, we first classified human protein-coding genes into 26 evolutionary age categories according to the gene age annotation framework proposed by Yin et al^14^. This framework assigns each gene to the phylogenetic node at which its orthologs first appear, spanning from the origin of life (∼4 billion years ago) to the present. Our analysis revealed that more than 90% of CDGs originated by the Devonian period (∼492 million years ago) or earlier (Figure 2A; Supplementary Table 2), suggesting that CDGs predominantly constitute evolutionarily conserved components of the core genomic machinery essential for multicellular life. Gene Ontology (GO) enrichment analysis further showed that these genes are preferentially associated with ancient core modules underlying fundamental biological processes, including cellular homeostasis, regulation of cell proliferation, and tissue regeneration (Supplementary Figure 3).

**Figure 2.**
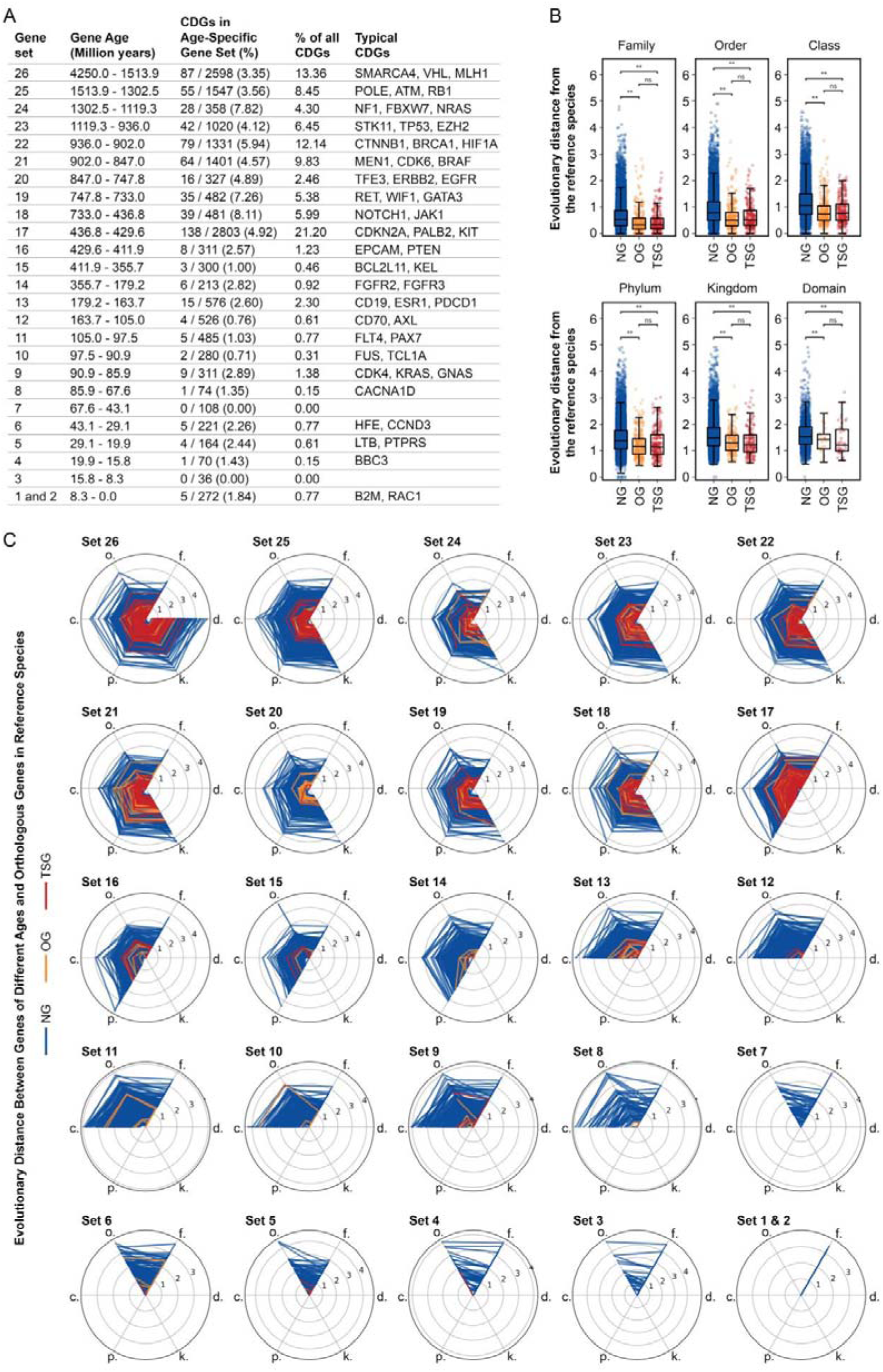
Comparison of structural semantic constraints between CDGs and non-CDGs across species evolution. A: Distribution of CDGs based on their evolutionary age (time of first appearance in the phylogenetic tree). B: Comparison of SSED distances between non-CDGs, OGs, and TSGs and their corresponding last common ancestors across taxonomic levels from Kingdom to Genus (Mann-Whitney U test). C: Comparison of SSED distances to the last common ancestor between non-CDGs and CDGs across different gene age strata. d: Domain, k.: Kingdom, p: Phylum, c.: Class, o.: Order, f.: Family, ns: No significant difference, *: P<0.05, **: P<0.005.

We then compared the SSED distances from human OGs, TSGs, and non-CDGs to their most recent common ancestors across multiple taxonomic levels (from kingdom to genus). Both OGs and TSGs exhibited significantly lower SSED values than non-CDGs across all taxonomic levels (Figure 2B), whereas no significant differences were observed between OGs and TSGs. Together, these results indicate that CDGs are subject to stronger evolutionary constraints, which maintain their structural and functional stability by restricting the range of structural–semantic changes tolerated during evolution.

To assess the widespread nature of these evolutionary constraints across different gene ages, we stratified genes by their origin time and compared the SSED values of CDGs and non-CDGs across various evolutionary periods. CDGs consistently exhibited lower SSED values, regardless of their origin in prokaryotic/early eukaryotic periods or more recently within the mammalian lineage, across various taxonomic levels (Figure 2C). These findings show that the evolutionary constraints on CDGs are not only linked to their early origins but also reflect widespread and persistent structural semantic constraints throughout evolution.

### 3 Oncogenic mutations conform to evolutionary constraints

Under the hypothesis of evolutionary constraints, a natural question arises: If the structural semantics of CDGs are widely constrained by evolutionary mechanisms, do the oncogenic mutations in CDGs face the same constraints? To test this, we focused on Class I oncogenic mutations (those with clear clinical and biological evidence linking their alterations to malignancies) from the COSMIC database (Supplementary Table 3). We constructed random mutation sets for the same genes and compared the SSED values between Class I oncogenic mutations and random mutations relative to their corresponding wild-type genes. The results showed that in almost all genes, the SSED between Class I mutations and the wild-type gene was significantly smaller than that of random mutations, suggesting a stronger evolutionary constraint on oncogenic mutations (Figure 3A). To rule out the potential impact of mutation sites on structural changes, we further constructed sets of site-matched control mutations (mutations at the same sites but with different amino acid changes). Under these conditions, Class I mutations still exhibited significantly smaller SSED values than these controls (Figure 3B). These results indicate that oncogenic mutations cause smaller structural semantic alterations, consistent with the hypothesis that CDGs are under stronger evolutionary constraints.

**Figure 3.**
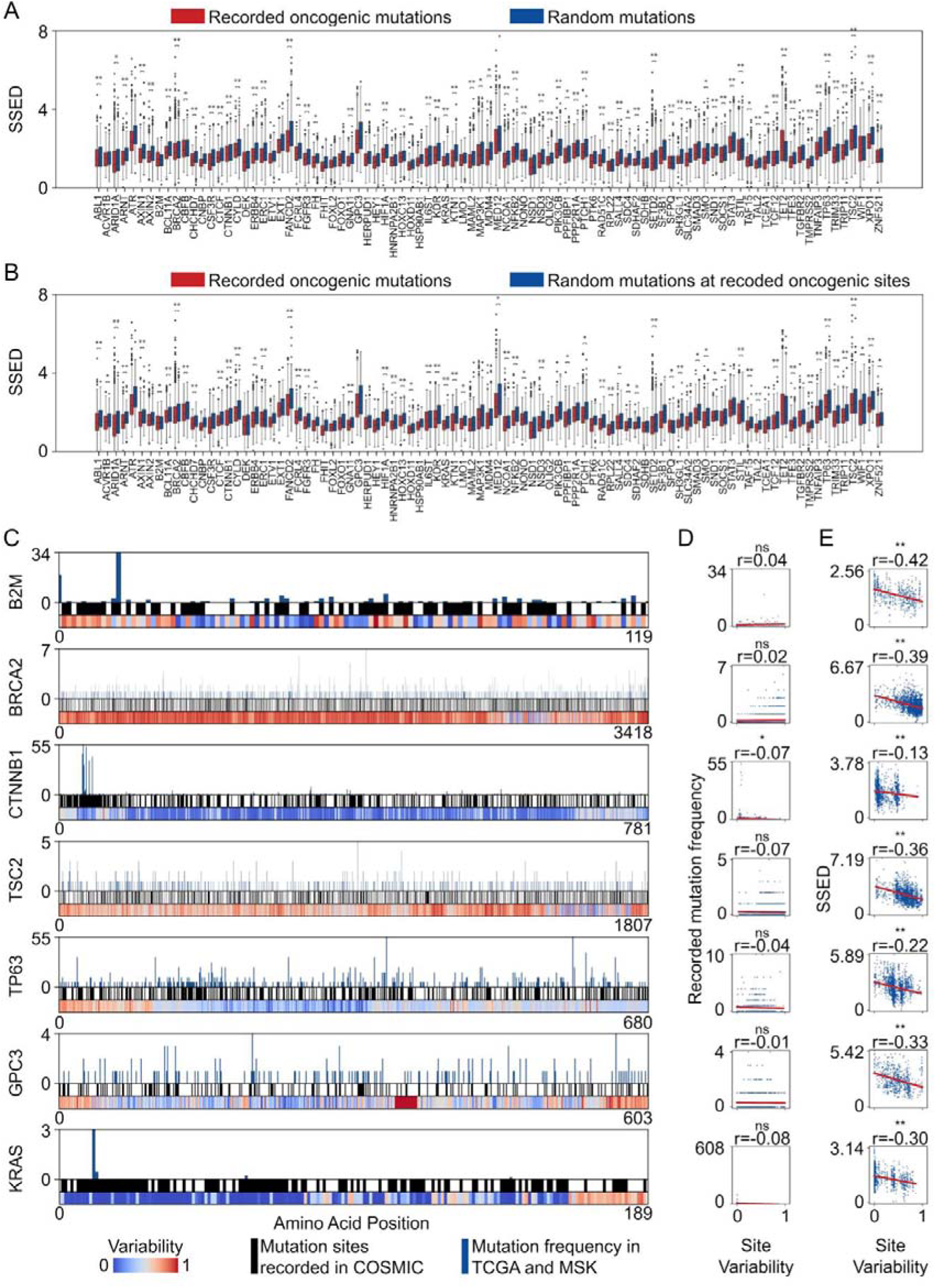
Oncogenic mutations are subject to stronger structural semantic evolutionary constraints. A: Comparison of SSED distances to the wild-type protein for documented oncogenic mutations vs. random mutations in the same genes, for Class I CDGs recorded in the COSMIC database. B: Comparison of SSED distances between Class I oncogenic mutations and other amino acid substitutions at the same sites (Mann-Whitney U test). C: For representative Class I CDGs, the evolutionary variability at each amino acid site is plotted against documented mutation sites (COSMIC) and their mutation frequencies (TCGA and MSK). D: Spearman’s rank correlation analysis between amino acid site evolutionary variability and recorded mutation frequency. E: Spearman’s rank correlation analysis between amino acid site evolutionary variability and the SSED distance of mutations at those sites. ns: No significant difference, *: P<0.05, **: P<0.005.

We next examined the relationship between site variability and oncogenic mutations. Site variability was quantified as the normalized pairwise Hamming distance between amino acid residues across 18,246 reference species with homologous sequences (Supplementary Table 4), providing a measure of evolutionary conservation. For representative genes from different evolutionary age categories, we plotted the variability of each site against the documented mutation sites and their frequencies from the TCGA database. No significant correlation was observed between site variability and the recorded mutation frequencies (Figure 3C, D). However, further analysis revealed a significant negative correlation between site variability and the SSED of mutations at those sites in most genes (Figure 3E). This indicates that sites with lower variability (i.e., more evolutionarily conserved) are associated with larger SSEDs when mutated.

### 4 Minimal structural semantic perturbations drive clonal selection and tumorigenesis

To investigate the relationship between the SSED of CDG mutations and clonal expansion capacity, we established stable cell lines expressing four different KRAS mutants (R135T, R135L, R135N, R135P) at residue 135. These mutants were introduced into two normal human epithelial cell lines: human pancreatic ductal epithelial cells (hTERT-HPNE) and human bronchial epithelial cells (BEAS-2B) (Supplementary Table 5). These mutants exhibited varying SSED values compared to the wild-type (WT) protein: R135T (0.7), R135L (1.2), R135N (8.0), and R135P (11.2) (Figures 4A, B; Supplementary Figure 4). Among these, R135T is a known oncogenic mutation, whereas the other mutations (R135L, R135N, and R135P) were previously uncharacterized.

**Figure 4.**
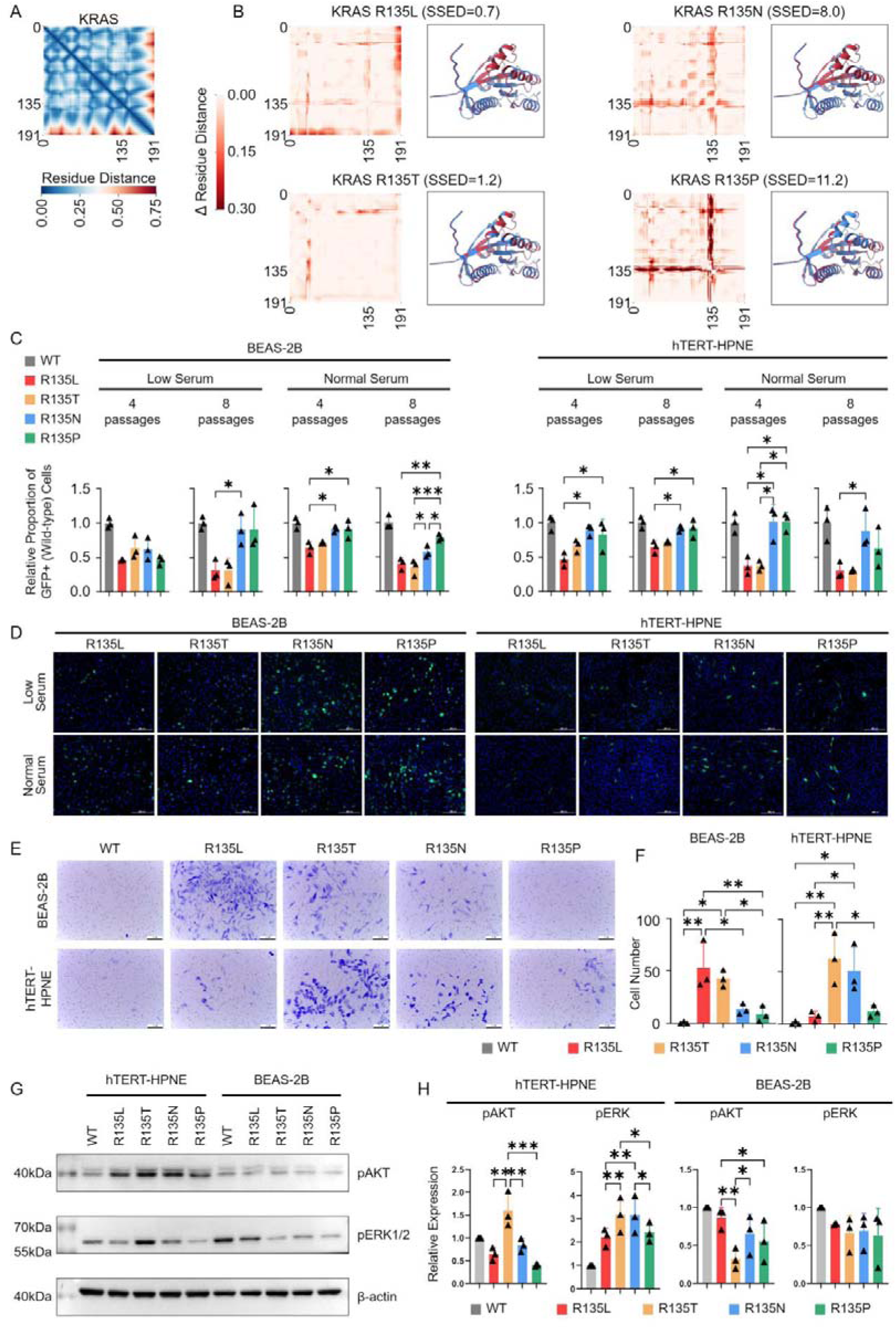
Association between the SSED of mutations and cellular proliferation, invasion, and signaling pathway activation. A: Amino acid residue contact matrix of the wild-type KRAS protein. B: Differences in the residue contact matrix and superimposed 3D structural (predicted by ESM3) comparisons between the wild-type KRAS protein and four mutants (R135T, R135L, R135N, and R135P). C: Clonal competition assay assessing the proliferative advantage of mutants with different SSED values. Quantitative analysis (mean±SEM, n = 3, one-way ANOVA) of the proportion of GFP-positive cells (representing the wild-type population) after co-culture with mutants in BEAS-2B and hTERT-HPNE cell lines under normal (10% FBS) and low-nutrient stress (0.5% FBS) conditions. D: Representative fluorescence images of the co-culture (scale bar: 200μm). E, F: Transwell invasion assay evaluating the invasive capacity of different mutants. E: Representative images of invaded cells. F: Quantitative analysis of invaded cells (mean±SEM, n = 3, one-way ANOVA). G, H: Western blot analysis of downstream signaling pathway activation. G: Representative bands for p-AKT, p-ERK1/2, and β-actin. H: Quantitative analysis of pAKT and pERK1/2 (normalized to total protein and β-actin, mean±SEM, n = 3, one-way ANOVA). WT: wile-type, ns: No significant difference, *: P<0.05, **: P<0.005, ***: P<0.0005.

First, we assessed the proliferative advantage using a competitive co-culture assay. When mutant cells were co-cultured with GFP-labeled WT cells, the proportion of cells harboring lower SSED mutations (R135L/R135T) increased significantly compared to cells with higher SSED mutations (R135N/R135P) after both 4 and 8 passages in both hTERT-HPNE and BEAS-2B lines. This proliferative advantage was consistent under both normal and low-serum (0.5% FBS) stress conditions (Figures 4C, D), indicating that lower SSED mutations confer a stronger clonal expansion capacity.

Subsequently, Transwell invasion assays revealed that in BEAS-2B cells, both lower SSED mutants (R135L and R135T) exhibited significantly increased invasiveness and migration compared to higher SSED mutants and WT cells. In hTERT-HPNE cells, the R135T mutant showed the strongest invasiveness, while the R135L mutant displayed no significant advantage. The R135N mutant demonstrated relatively high invasiveness in this cell line (Figures 4E, F). Analysis of key KRAS downstream oncogenic signaling pathways by Western blot showed that in BEAS-2B cells, the R135L mutant had the highest pAKT levels, while pERK1/2 levels showed no significant differences among mutants. In hTERT-HPNE cells, the R135T mutant exhibited the highest pAKT and pERK1/2 levels, whereas the R135N mutant showed elevated pERK1/2 but not pAKT levels (Figures 4G, H). Although some cell-type specificity was observed, the overall trend indicates that lower SSED mutations are associated with stronger oncogenic phenotypes and signaling pathway activation.

To validate the in vivo relevance of these findings, we conducted tumor formation assays using nude mice. Three weeks after inoculation, stable tumors formed in three mice (Figure 5A), all originating from BEAS-2B cells (two with the R135L mutation and one with R135T) (Figure 5B). Five additional mice developed transient visible tumors (including one BEAS-2B R135L, two BEAS-2B R135T, and two hTERT-HPNE R135L), which later regressed (Figure 5C). Histological analysis of the tumors confirmed their malignant characteristics. H&E staining revealed nuclear enlargement, hyperchromasia, and an increased nuclear-to-cytoplasmic ratio, while immunohistochemistry confirmed CK7 positivity and Ki-67 expression (Figure 5D). These results validate KRAS R135L as a previously unreported oncogenic mutation in bronchial epithelial cells within the “evolutionary constraint” framework.

**Figure 5.**
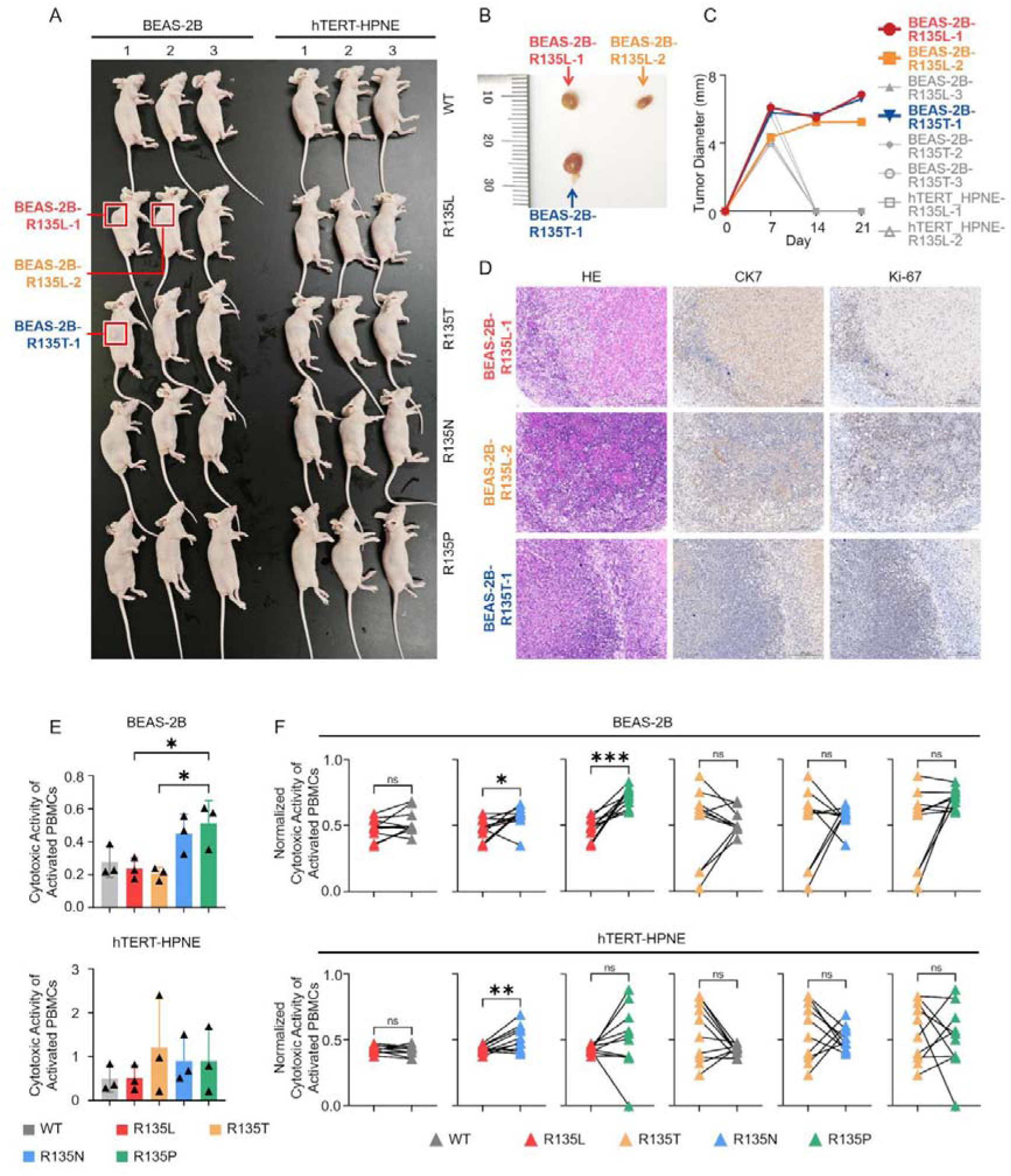
In vivo tumorigenicity and in vitro immunogenicity of KRAS mutants with different SSED values. A: Photographs of nude mice three weeks after inoculation with wild-type and four KRAS mutant BEAS-2B and hTERT-HPNE cells. Sites forming stable tumors are marked with red boxes. B: Photographs of stable tumors excised from the mice. C: Tumor growth curves. All observed tumors were recorded; stably formed tumors are shown in color, and regressed tumors are shown in gray. D: Histological analysis of tumor tissues. H&E staining shows tumor cell morphology; immunohistochemical staining detects expression of CK7 (epithelial marker) and Ki-67 (proliferation marker) (scale bar: 200μm). E, F: In vitro cytotoxicity of PBMCs against different mutant cells measured by LDH release assay. E: Cytotoxicity of pooled PBMCs (from 10 healthy donors) against BEAS-2B and hTERT-HPNE mutant cells (mean ± SEM, n=3, one-way ANOVA). F: Paired analysis based on individual donors, comparing the killing efficacy of PBMCs against high-versus low-SSED mutants (mean ± SEM, n=10, paired t-test). ns: No significant difference, *: P<0.05, **: P<0.005, ***: P<0.0005.

### 5 Immune recognition efficacy is governed by structural semantic distance under evolutionary constraint

To determine the impact of SSED on tumor cell immunogenicity, we assessed how different KRAS mutants affect immune-mediated killing. We co-cultured mutant hTERT-HPNE and BEAS-2B cell lines with pooled peripheral blood mononuclear cells (PBMCs) from 10 healthy donors and measuring cytotoxicity via LDH release. In BEAS-2B cells, the high-SSED mutant (KRAS R135P) was killed significantly more effectively than the low-SSED mutants (R135L and R135T). However, the killing efficacy of pooled PBMCs did not differ significantly among KRAS mutants with different SSED values in hTERT-HPNE cells (Figure 5E).

Since heterogeneity among donor PBMCs (e.g., in lymphocyte subsets and HLA genotypes) could mask immunogenicity differences against specific mutants, we performed a paired analysis at the individual donor level. This paired analysis revealed that, within the same donor’s PBMCs, the killing rate of the low-SSED mutant (KRAS R135L) in BEAS-2B cells was significantly lower than that of high-SSED mutants (R135N and R135P). Similarly, in hTERT-HPNE cells, the high-SSED mutant (R135N) was killed more efficiently than the low-SSED mutant (R135L) (Figure 5F). These results collectively suggest that mutations inducing larger structural semantic perturbations (high SSED) enhance immune recognition and killing. Conversely, mutants with minimal alterations (low SSED) are prone to immune evasion, thereby conferring a survival advantage under immune pressure.

### 6 The Impact of SSED on the efficacy of immune therapy

Based on cellular experiments that confirmed the association between SSED and immune cytotoxicity, we next explored its potential to predict the efficacy of immune checkpoint blockades (ICBs) in clinical settings. We hypothesized that tumors with mutations inducing higher SSED may generate more immunogenic neoantigens, thereby eliciting a stronger anti-tumor immune response. To test this hypothesis, we introduced the SSEDmax metric, which is defined as the highest SSED value across all mutations in a patient. This metric quantifies the maximum potential immunogenicity inherent in the patient’s tumor.

We conducted an external validation in a real-world cohort of 576 lung cancer patients from the MSK center who were treated with ICBs. Survival analysis showed that patients with high SSEDmax had significantly longer progression-free survival (PFS) and overall survival (OS) compared to those with low SSEDmax, regardless of whether ICBs were used as first-line or subsequent-line therapy (Figure 6A). SSEDmax showed no significant correlation with clinical features such as age, sex, or AJCC stage (Figure 6B-D). Notably, SSEDmax was positively correlated with tumor mutational burden (TMB) (Figure 6E), suggesting that tumors with higher mutation loads are more likely to generate highly immunogenic mutations.

**Figure 6.**
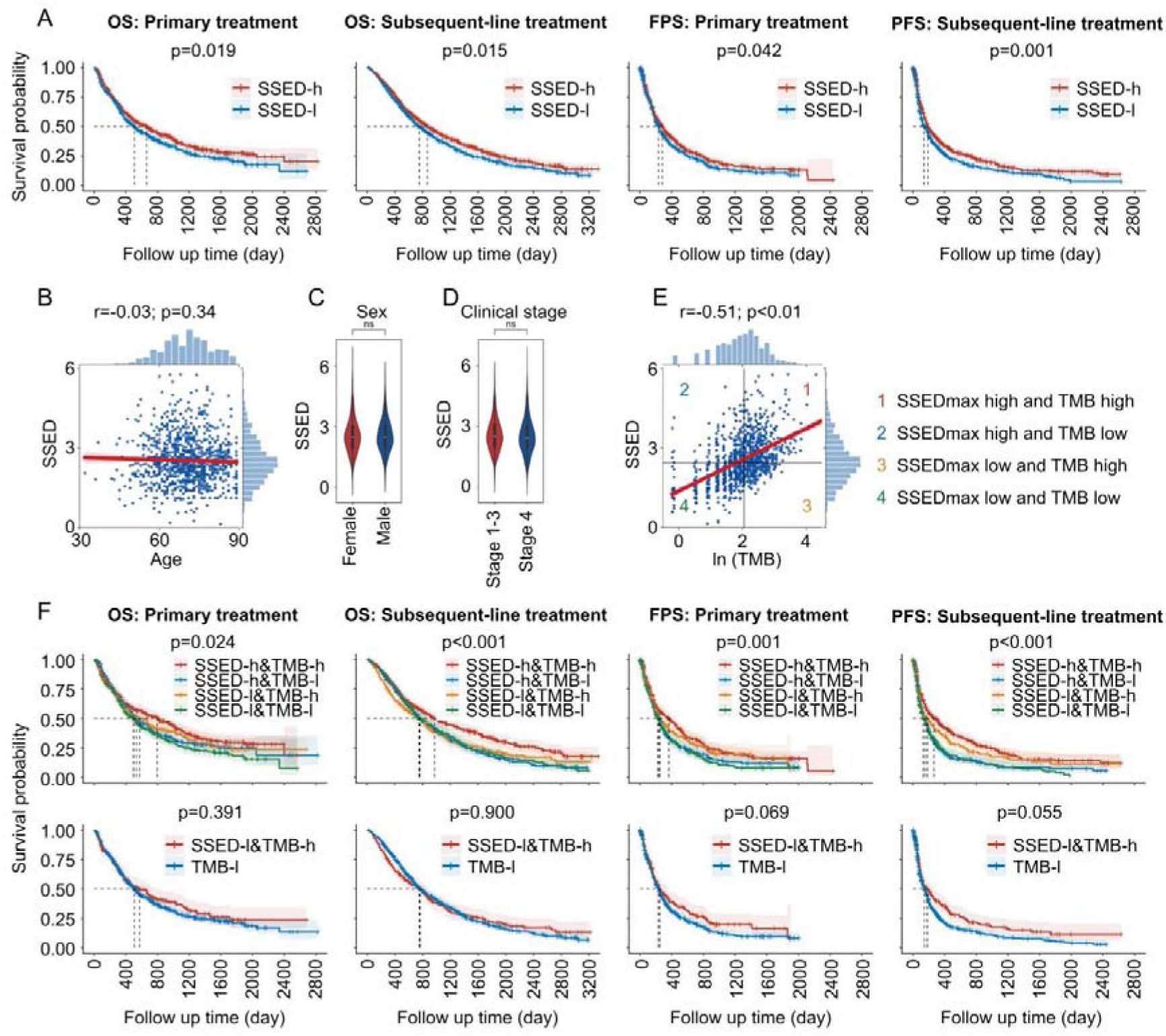
Correlation analysis of SSEDmax with clinical features and efficacy of immune checkpoint inhibitor therapy in lung cancer patients (MSK cohort). A: PFS and OS for patients with high vs. low SSEDmax receiving ICI therapy (Kaplan-Meier curves, Log-rank test). B: Spearman correlation analysis between SSEDmax and patient age. C: Comparison of SSEDmax between sexes (t-test). D: Comparison of SSEDmax between AJCC stages (I–III vs. IV) (t-test). E: Spearman correlation analysis between SSEDmax and TMB. F: Survival analysis based on SSEDmax and TMB (Kaplan-Meier curves, Log-rank test). (Top) PFS and OS for four patient subgroups: SSEDmax-high/TMB-high, SSEDmax-high/TMB-low, SSEDmax-low/TMB-high, and SSEDmax-low/TMB-low. (Bottom) Comparison of PFS and OS between TMB-high/SSEDmax-low patients (excluding SSEDmax-high patients from the TMB-high group) and TMB-low patients. SSED-h: SSEDmax high, SSED-l: SSEDmax low, TMB-h: TMB high, TMB-l: TMB low.

To investigate the relationship between SSEDmax and TMB, and their combined impact on prognosis, we stratified patients into four groups based on the median values of both metrics: SSEDmax-high/TMB-high, SSEDmax-high/TMB-low, SSEDmax-low/TMB-high, and SSEDmax-low/TMB-low. Survival analysis revealed that only the SSEDmax-high/TMB-high group showed a significant advantage in PFS and OS. Importantly, among the TMB-high subgroup, patients with low SSEDmax (TMB-high/SSEDmax-low) showed survival benefits comparable to those with TMB-low, indicating that high TMB alone does not provide a significant advantage when excluding SSEDmax-high patients (Figure 6F). These results establish SSEDmax as an effective biomarker for predicting ICI response, independent of standard clinical features and complementary to TMB. While TMB quantifies the quantity of neoantigens, SSEDmax captures the immunogenic quality of the most divergent neoantigen. These two metrics represent orthogonal dimensions for assessment, providing a more robust foundation for predicting treatment response and refining patient selection for immunotherapy.

## Discussion

Our findings reveal that CDGs are subject to stringent evolutionary constraints, a phenomenon consistently observed across CDGs from different evolutionary time points. Moreover, oncogenic mutations adhere to these same constraints, suggesting they are subject to evolutionary pressures. These results provide a unified explanation for the evolutionary selection patterns of CDGs, spanning both long evolutionary periods (hundreds of millions of years) and shorter time frames (tumorigenesis at the individual level).

To incorporate both structural and functional information into evolutionary distance assessments, we propose a novel approach (SSED), which is based on protein language models. Compared with sequence analysis, protein function is more closely tied to its three-dimensional structure, making a structure-based evolutionary distance metric a more effective reflection of the functional selection pressures proteins undergo in evolution. By leveraging the ESM3 protein language model, which utilizes large datasets of protein structures, functional annotations, and homologous sequence alignments, SSED incorporates additional evolutionary insights, enhancing its accuracy. Notably, the ESM3 model can generate proteins with only 58% sequence similarity to existing proteins but with similar functions, demonstrating its ability to capture structural and functional nuances^13^. Our results show that SSED provides a more accurate measure of evolutionary distance between proteins and helps construct reliable phylogenetic relationships.

From an evolutionary perspective, many CDGs have ancient origins, with a concentration in early eukaryotic and pre-metazoan stages, supporting previous studies^14,15^. Functionally, many CDGs are crucial for core cellular processes, including the cell cycle, DNA repair, and apoptosis inhibition^16–18^. On one hand, changes in these proteins must be tightly controlled to preserve their core functionality, while on the other hand, they must undergo modifications under evolutionary pressure to meet the demands of multicellular organisms^19,20^. The balance between structural stability and evolutionary adaptability likely explains the stronger constraints on the structural semantics of CDGs.

Regarding the evolutionary role of oncogenic mutations, some studies suggest that they follow a “degenerative” pattern, where cancer cells progressively shed multicellular constraints, resembling a reversion to a unicellular ancestral state^15,21^. Conversely, other studies argue that oncogenic mutations are not degenerative but instead have adaptive significance. Research by McFarland et al.^22^highlights that driver mutations provide proliferative advantages, indicating that oncogenic mutations promote the adaptive evolution of malignant cells. A study by Gu et al.^23^ suggests that 35 CDGs, spanning 47 types of malignant tumors, are under positive selection in modern human populations. Our study supports the view that oncogenic mutations are not deleterious events, but rather extensions of the long-term evolutionary constraints of CDGs. Oncogenic mutations require sufficient functional alterations to gain proliferative advantages, but these changes must remain within limits to avoid immune destruction. The evolutionary constraint mechanism defines the structural semantic range of oncogenic mutations, balancing clonal expansion and immune evasion, and granting oncogenic mutations an intrinsic evolutionary drive.

In functional cell experiments, our results support the hypothesis of a dual constraint model, where low SSED (proximal mutations) maintain protein function and escape immune recognition, whereas high SSED (distal mutations) are likely cleared by immune responses due to stronger immune activation. This phenomenon further supports the notion that oncogenic mutations result from a balance between clonal expansion and immune evasion. Importantly, this suggests that SSED reflects immune response strength and offers a novel metric for predicting immune therapy efficacy, as well as for selecting patient populations for immunotherapy. In clinical data, we found that SSEDmax significantly influences PFS and OS during immune therapy, independent of traditional TMB metrics. Although TMB measures mutation count, it does not assess whether these mutations can trigger sufficient immune responses. As a supplement, SSED quantifies the extent of structural semantic changes induced by mutations, reflecting whether these changes are sufficiently recognized by the immune system. Therefore, TMB and SSED represent two orthogonal dimensions: TMB quantifies the number of neoantigens, while SSED evaluates the quality of neoantigens relative to wild-type genes. This composite model of antigen quantity and quality provides a new explanation for clinical heterogeneity in immune therapy, such as the lack of response in TMB-high patients and the clinical benefit in TMB-low patients. SSED shows promise as a valuable supplementary indicator for predicting immune therapy efficacy, offering new theoretical insights into immune response heterogeneity and optimizing patient selection strategies.

This study has several limitations. First, to ensure result comparability, we focused solely on single-site amino acid mutations, though CDGs can also undergo alterations such as amplifications, truncations, and insertions. A unified computational framework to quantify the evolutionary distance changes reflected by these alterations is still needed. Furthermore, even identical mutations at the same site of the same gene (e.g., KRAS R135T in this study) appear to utilize distinct pathways (such as AKT or ERK) in different cell types to enhance adaptability and invasiveness. The underlying mechanisms responsible for this phenomenon remain to be elucidated.

In conclusion, this study introduces and validates the concept of “evolutionary constraint” in CDGs, establishing a framework that bridges long-term species evolution with short-term tumor adaptation. This framework offers theoretical insights and novel quantitative tools for patient stratification and efficacy prediction in immune therapy.

## Materials and Methods

### Calculation of SSED

The structural semantic representations of amino acid sequences were extracted with the pre-trained ESM-3 protein language model (version 3.0.1). Each amino acid sequence was processed through the ESM-3 encoder and decoder, and the resulting structure tokens—representing the learned structural semantic features—were extracted. Multiple sequence alignment was done with MAFFT (NCBI BLAST+ version 2.13.0, default parameters), and residues were aligned to establish positional correspondence between sequences. The structural evolutionary distance between any two homologous protein sequences was defined as the average Minkowski distance (p=2) between all aligned structure token pairs. Specifically, let 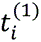 and 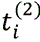 denote the structure tokens at the i-th aligned amino acid position for the two sequences. The SSED distance *D* is calculated as:

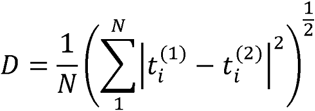

where *N* represents the number of valid aligned positions. The SSED calculation code used in this study is publicly available at https://github.com/FreudDolce/SSED/.

### Benchmarking and phylogenetic tree reconstruction

To evaluate the performance of SSED in reflecting phylogenetic relationships, we generated datasets containing between 1 and 3000 protein-coding genes (total amino acid length range: 15,652–5,315,823) from representative species and calculated pairwise SSED distances between all species to construct distance matrices. Phylogenetic trees were reconstructed from these matrices using the neighbor-joining method. Amino acid sequences for each species were obtained from the OrthoDB database (v11). Three representative taxonomic groups—Mammalia, Thermoproteota, and Rhodobacteraceae—were selected to evaluate SSED-based phylogenetic reconstruction across different evolutionary scales. For subsequent analyses involving CDGs, a standard list of CDGs was compiled from the OncoKB database (data retrieved December 2024). CDGs were classified as oncogenes or tumor suppressor genes according to their functional annotations. Non-CDGs were defined as protein-coding genes not annotated as CDGs in OncoKB.

The NJ and UPGMA, using the BLOSUM62 amino acid substitution model, were used as sequence-based controls. Both are widely used methods for phylogenetic tree construction based on amino acid sequences. These methods were implemented using the Python Phylip library, with UPGMA assuming a constant evolutionary rate. Phylogenetic trees from TimeTree.org (data retrieved April 2025) were used as reference phylogenies. Topological similarity between inferred and reference trees was assessed using the unweighted Robinson–Foulds distance.

### Gene age annotation and phylogenetic stratification

To investigate the evolutionary origins of CDGs, we applied the gene age annotation method proposed by Yin et al.^14^. This method classifies each human protein-coding gene into one of 26 origin-time categories based on its first appearance in the species phylogenetic tree, covering 4 billion years of evolutionary history, from the origin of life to major branching events like the last universal common ancestor (LUCA). Briefly, the method begins with a whole-genome homology search using BLASTP, followed by homologous gene clustering with the Markov Cluster Algorithm (MCL). Phylogenetic inference is then used to construct a maximum likelihood evolutionary tree, integrating both horizontal gene transfer and vertical evolution to pinpoint gene duplication nodes. This process assigns human genes into 26 refined age categories.

### Calculation of amino acid site variability

In this study, we defined the evolutionary variability of a site based on normalized pairwise Hamming diversity, which represents the average divergence at that site across a multiple sequence alignment of homologous proteins from multiple species. For each site, we performed pairwise comparisons of amino acid residues from homologous sequences across 18,246 reference species. A comparison was scored as 1 if the two residues differed, and 0 otherwise. The sum of all pairwise mismatches was calculated and normalized to a variability score using the following formula:

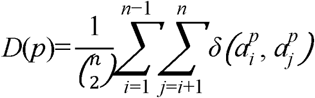

where *D*(*p*) denotes the variability at position *p*; *n* is the number of homologous proteins from reference species; 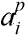 or 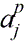 represents the amino acid at position *p* for the *i^th^* or *j^th^* species, respectively; δ(x, y) is the indicator function, which equals 1 if *x*≠*y* and 0 otherwise; and 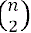 is the total number of comparable species pairs.

### Cell lines and culture conditions

This study used two human immortalized epithelial cell lines as model systems for functional experiments: the human pancreatic ductal epithelial cell line hTERT-HPNE and the human bronchial epithelial cell line BEAS-2B. All cell lines were purchased from the American Type Culture Collection (ATCC) and cultured according to the recommended conditions at 37°C in a humidified atmosphere containing 5% CO[. Both cell lines were maintained in Dulbecco’s Modified Eagle Medium (DMEM, Gibco, C11995500BT) supplemented with 10% fetal bovine serum (FBS, ExCell, FSP500). All experiments were performed using cells within 20 passages to ensure consistency. For stress simulation assays, cells were cultured in low-serum medium containing 0.5% FBS, while all other culture parameters were identical to the standard conditions.

### Construction of stable mutant cell lines

To establish stable cell lines expressing different KRAS mutants, we used human KRAS cDNA as a template and introduced four distinct mutations at the R135 residue of KRAS by site-directed mutagenesis. The resulting mutants were R135T, R135L, R135N, and R135P, with SSED values of 0.7, 1.2, 8.0, and 11.2, respectively. The mutant sequences were cloned into the pLenti-SFFV-Puro lentiviral vector. Viral particles were packaged in HEK293T cells by co-transfecting the transfer plasmid with the packaging plasmids psPAX2 and pMD2.G. Seventy-two hours post-infection of the target cells (hTERT-HPNE and BEAS-2B), selection was initiated with 2μg/mL puromycin to obtain polyclonal populations stably expressing the KRAS mutants. Successful viral transduction and genomic integration were confirmed by PCR amplification of the transgene, and KRAS (Flag) protein expression levels were verified by Western blot analysis.

### Detection of RAS downstream signaling

To assess the impact of different KRAS mutants on activation of the RAS downstream signaling axis, we measured the levels of phosphorylated ERK (pERK) and phosphorylated AKT (pAKT) by Western blot analysis. Proteins were separated by SDS–PAGE and transferred onto membranes. Membranes were incubated with antibodies against pERK (Thr202/Tyr204, Proteintech, 28733-1-AP), pAKT (Ser473, Proteintech, 66444-1-Ig), and β-actin. Signal detection was performed using HRP-conjugated secondary antibodies and ECL chemiluminescence. Band densities were analyzed with ImageJ software and normalized to the corresponding total pERK or pAKT levels and to β-actin.

### Cell invasion assay

Cell invasion capacity was assessed using Transwell chambers (Corning, 8.0 μm pore size). Prior to the assay, the upper surface of the polycarbonate membrane was coated with 50 μL of Matrigel matrix (Corning, #356234, diluted 1:8 in serum-free DMEM) and incubated at 37°C for 2 hours to allow polymerization. Cells (wild-type and KRAS mutant BEAS-2B and hTERT-HPNE cells) in the logarithmic growth phase were harvested, resuspended in serum-free medium, and adjusted to a density of 1.0×10□ cells/mL. A 100μL aliquot of the cell suspension (1.0×10□ cells per well) was added to the upper chamber, while the lower chamber was filled with 600μL of complete medium containing 20% FBS as a chemoattractant. Cells were allowed to invade for 24 hours at 37°C under 5% CO□. After incubation, non-invading cells on the upper surface of the membrane were gently removed with a cotton swab. Cells that had migrated to the lower surface were fixed with 4% paraformaldehyde for 15 minutes and stained with 0.1% crystal violet for 20 minutes. After rinsing with PBS, the membranes were imaged, and the number of invaded cells was counted in five randomly selected fields of view per membrane under a microscope. Each experimental group was performed in triplicate, and the entire experiment was independently repeated three times.

### Clonal competition assay

To compare the clonal advantage of cells expressing different KRAS mutants under competitive conditions, mutant cells were mixed with wild-type KRAS cells stably expressing GFP at a 1:1 ratio and co-cultured in 6-well plates. The co-culture was maintained under normal (10% FBS) and low-nutrient stress (0.5% FBS) conditions. Cells were harvested after 4 and 8 passages. The proportion of GFP-positive cells was determined by analyzing fluorescence images captured under a fluorescence microscope using ImageJ software. The clonal competitiveness of each mutant was evaluated by calculating the ratio of non-GFP (mutant) cells to the total population.

### Mouse xenograft tumor formation assay

Animal experiments were performed using 6–8-week-old female BALB/c nude mice. To monitor tumor formation and growth, each mouse received a single subcutaneous injection of a mutant cell line (1×10[cells in 100μL of a PBS/Matrigel mixture) into the right flank. Tumor dimensions were measured every 7 days. The experiment was terminated after 3 weeks or when the tumor volume exceeded 1500mm³. At the endpoint, mice were euthanized, and tumors were excised and weighed. All animal procedures were conducted in accordance with the Regulations for the Management of Laboratory Animals and were approved by the Institutional Animal Care and Use Committee of our institution (ethical approval protocol number: IACUC-20241034).

### Hematoxylin and Eosin (H&E) staining

Excised mouse tumor tissues were fixed in 4% paraformaldehyde for 24 hours, followed by conventional paraffin embedding and sectioning. Tissue sections were baked at 60°C for 1 hour, deparaffinized in xylene, and rehydrated through a graded series of ethanol. Sections were stained with hematoxylin for 5 minutes, rinsed in running tap water to develop the blue color, and differentiated in 1% acid alcohol for a few seconds. This was followed by counterstaining with eosin for 2 minutes. Sections were then dehydrated through graded ethanol, cleared in xylene, and mounted with neutral balsam. Images were acquired using a light microscope to evaluate the morphological characteristics of tumor cells, including nuclear size, hyperchromasia, nuclear-to-cytoplasmic ratio, and tissue architecture.

### Immunohistochemical (IHC) staining

After deparaffinization and rehydration (as described for H&E staining), paraffin sections were subjected to heat-mediated antigen retrieval in citrate buffer (pH 6.0). Endogenous peroxidase activity was quenched by incubating the sections with 3% hydrogen peroxide for 10 minutes at room temperature. Sections were then incubated with primary antibodies, including anti-cytokeratin 7 (CK7, Proteintech, 15539-1-AP) or anti-Ki-67 (Cell Signaling Technology, #9129T), at 4°C overnight. After washing with PBS, the sections were incubated with an HRP-labeled secondary antibody for 30 minutes at room temperature. Signal detection was performed using a DAB chromogenic kit, and nuclei were counterstained with hematoxylin. Finally, the sections were dehydrated, cleared, and mounted.

### PBMC cytotoxicity assay (LDH method)

Target cells were seeded in 96-well plates at a density of 1×10^4^ cells per well and cultured for 24 hours to allow adherence. Pre-activated PBMCs were added according to the experimental groups, and the co-culture was maintained for 24 hours. After co-culture, the plates were centrifuged at 700×g for 5 minutes at room temperature. Subsequently, 50μL of supernatant from each well was carefully transferred to a new 96-well plate. Using the LDH Cytotoxicity Detection Kit (Thermo Fisher, #EEA013), 100μL of lysis solution was added to the Maximum LDH release control wells. After incubation, 50μL of supernatant was similarly aspirated from these wells. Then, 50μL of the LDH substrate reaction solution was added to all wells containing supernatant. The plate was protected from light and incubated at 37°C for 20 minutes. Finally, the reaction was stopped by adding 100μL of stop solution per well, and the absorbance was immediately measured at 490nm (OD490) using a microplate reader, with a reference wavelength of 690nm for zero correction. The spontaneous release control consisted of target cells alone with an equal volume of complete medium. The maximum LDH release control consisted of target cells treated with lysis solution. The medium background control contained only complete medium. Cytotoxicity was calculated using the formula: % Cytotoxicity = [(Experimental Group OD – Spontaneous Release OD) / (Maximum Release OD – Spontaneous Release OD)]×100%. Each experimental group was performed in triplicate, and the entire experiment was independently repeated three times.

### Clinical data sources

The independent validation dataset used in this study was obtained from the Memorial Sloan Kettering Cancer Center (MSK) cohort of non-small cell lung cancer (NSCLC) patients receiving ICI treatment, who were selected for further analysis. The data included mutation annotations, clinical variables (e.g., age, sex, disease stage), Tumor Mutational Burden (TMB), and clinical treatment response information. All data handling complied with relevant data use regulations and institutional ethical guidelines.

### Comparative analysis and statistical tests

Continuous variables that followed a normal distribution are presented as mean±standard deviation, while those with a non-normal distribution are described using median and interquartile range (IQR). Categorical data are summarized as frequencies and percentages (%). For comparisons between two groups, an independent samples t-test was used for normally distributed data with homogeneous variances, and the Mann-Whitney U test for non-normally distributed data. For comparisons among more than two groups, one-way analysis of variance (ANOVA) was used. Correlation analyses were performed using Spearman’s rank correlation. Survival analysis was conducted using the Kaplan-Meier method to estimate PFS and OS, with between-group differences assessed by the Log-rank test. A two-sided p-value < 0.05 was considered statistically significant. Survival analysis and visualization were performed using the survival and survminer packages in R (v4.3.0). Other statistical analyses were conducted using Python libraries, including statsmodels, scikit-learn, and statannotations. Data visualization was implemented with seaborn and matplotlib packages in Python.

