## Supplemental Figures for "Structural semantic evolutionary distance (SSED) unifies the selection of cancer driver genes across macroevolution and tumorigenesis"

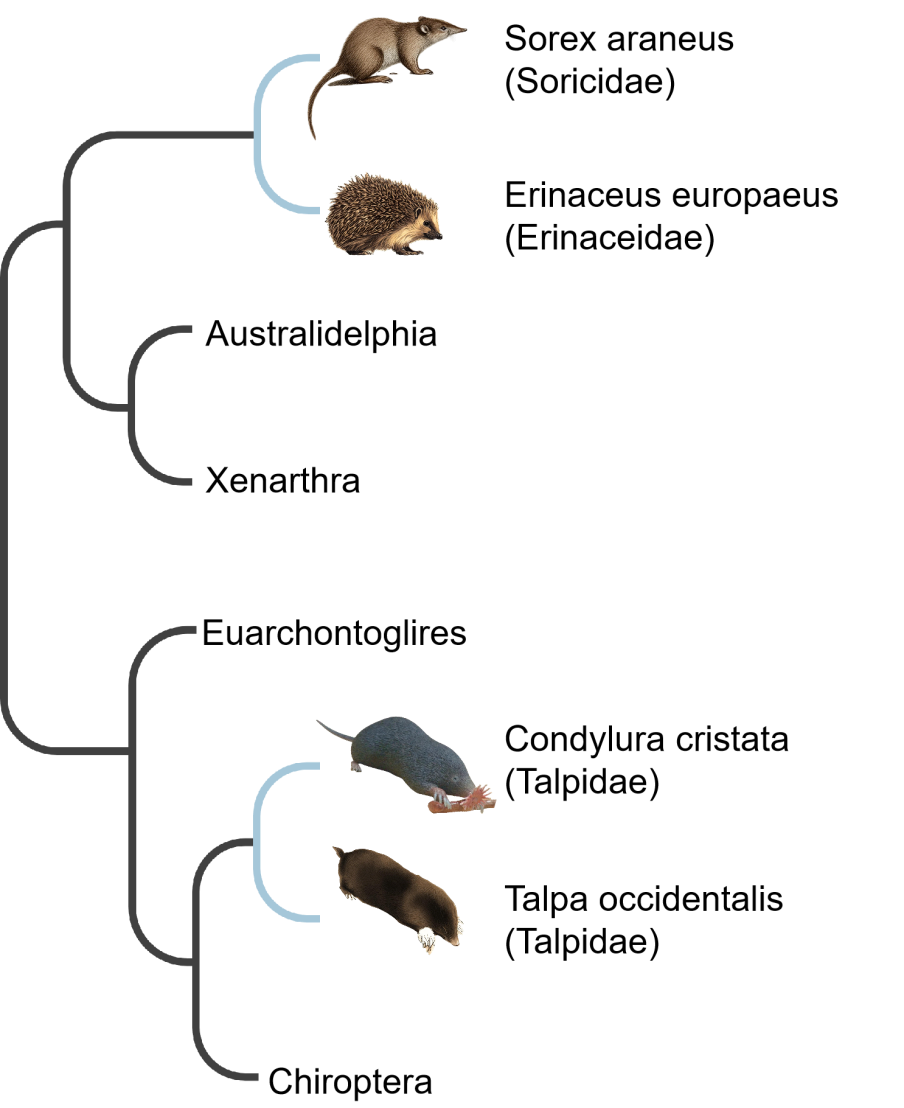

Supplementary Figure 1. The phylogenetic relationships within the order Eulipotyphla as resolved by the SSED tree.

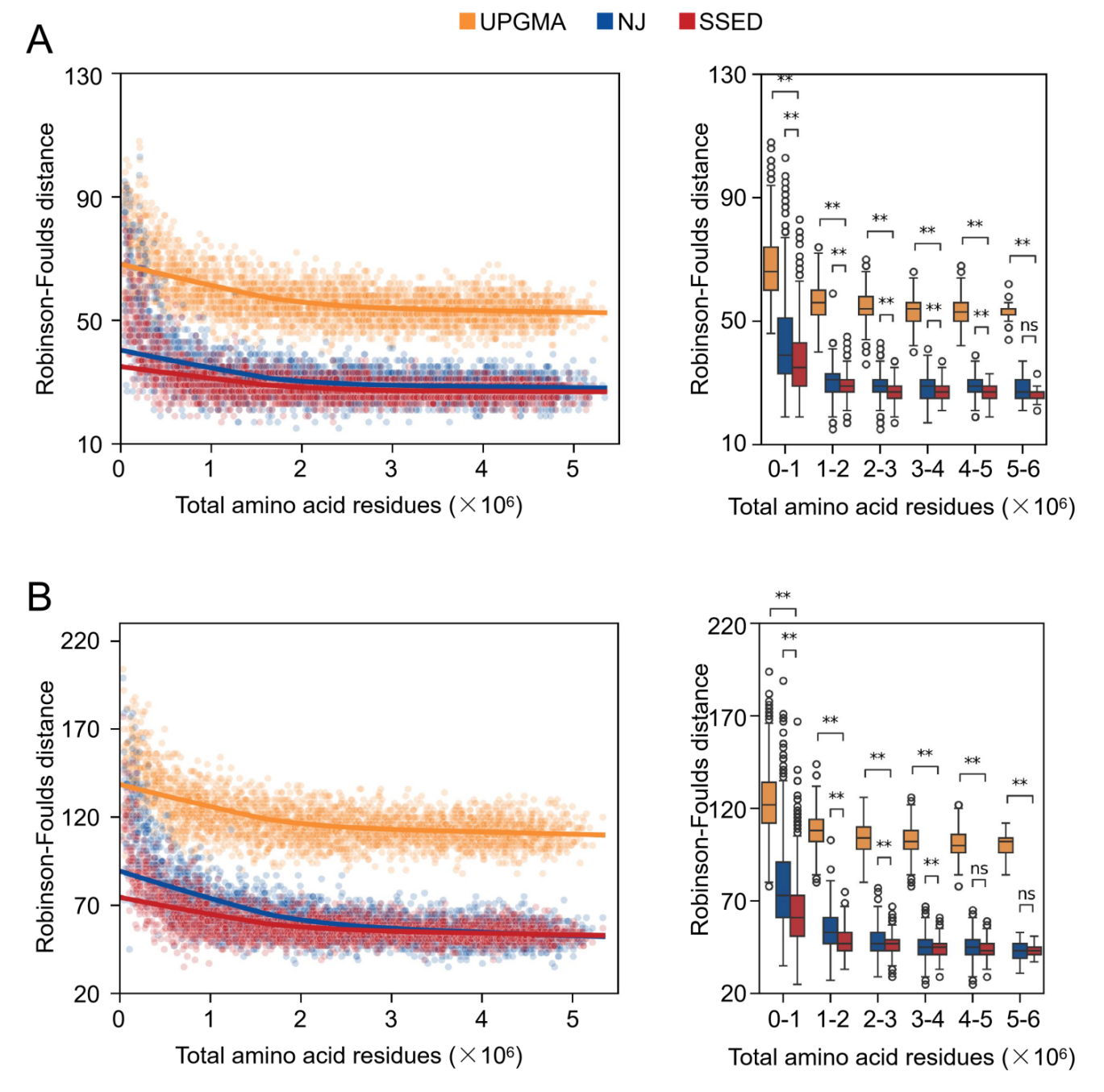

Supplementary Figure 2. Phylogenetic trees were inferred at two taxonomic levels: (A) the phylum Thermoproteota and (B) the family Rhodobacteraceae. The topological accuracy of trees built by SSED, neighbor-joining (NJ), and the unweighted pair group method with arithmetic mean (UPGMA) was assessed and compared using the unweighted Robinson-Foulds distance..

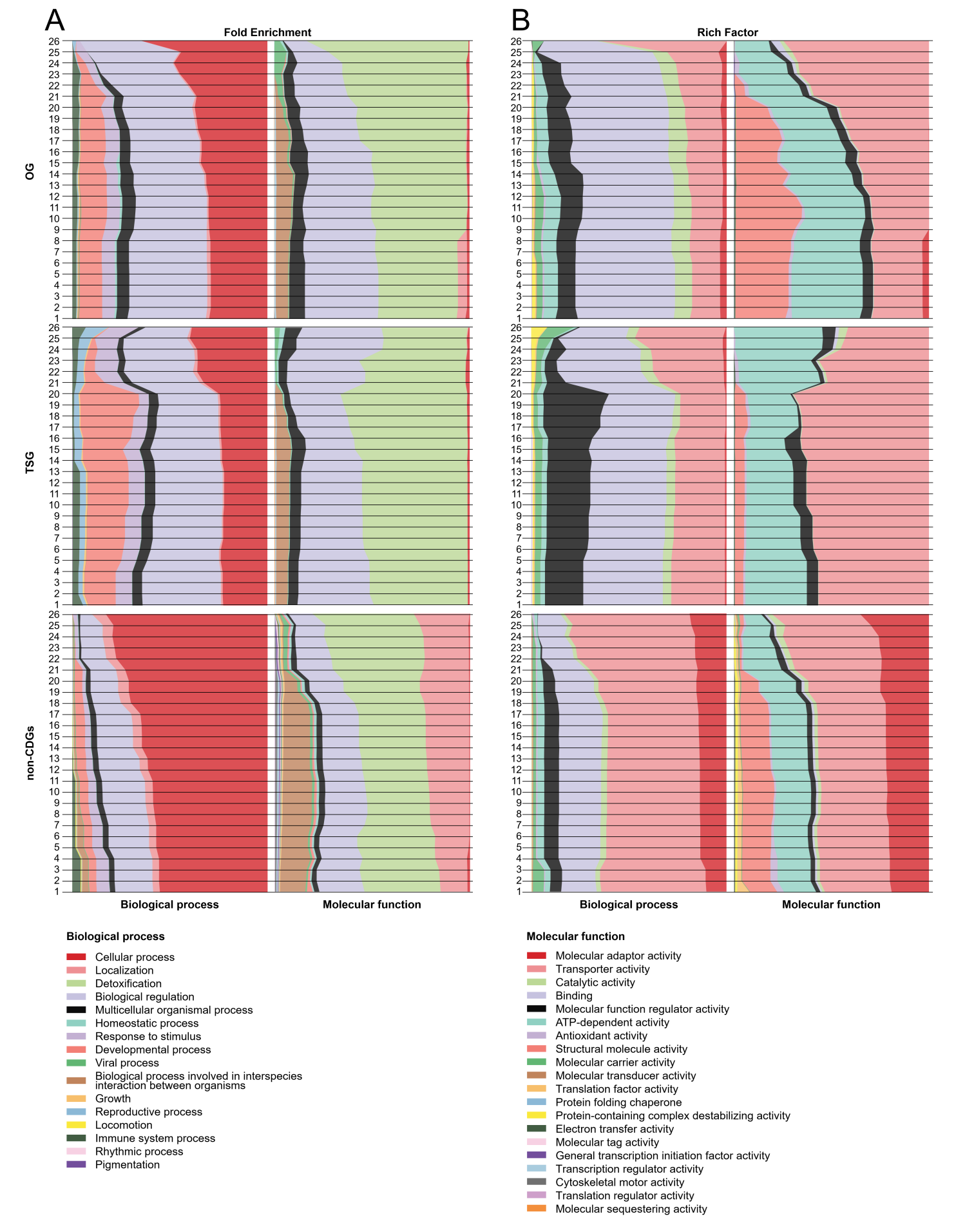

Supplementary Figure 3. Cumulative distribution and evolutionary trajectory of functional enrichment across age-stratified cancer driver genes. The changes in the functional enrichment profiles of OGs, TSGs, and non-CDGs as a function of cumulative gene age. Genes were stratified into 26 evolutionary age categories (from youngest, category 1, to oldest, category 26) based on their first appearance in the phylogenetic tree. A: Cumulative contribution of the gene sets to the Fold Enrichment in Gene Ontology (GO) terms. B: Cumulative contribution of the gene sets to the Rich Factor. For each gene category, moving from top to bottom, each bar represents the cumulative functional enrichment contribution from the specified age category inclusive of all older categories(e.g., the bar for age category 23 represents the cumulative contribution of genes in categories 23 through 26). All bars are normalized to show the relative proportion of contributions from different functional categories.

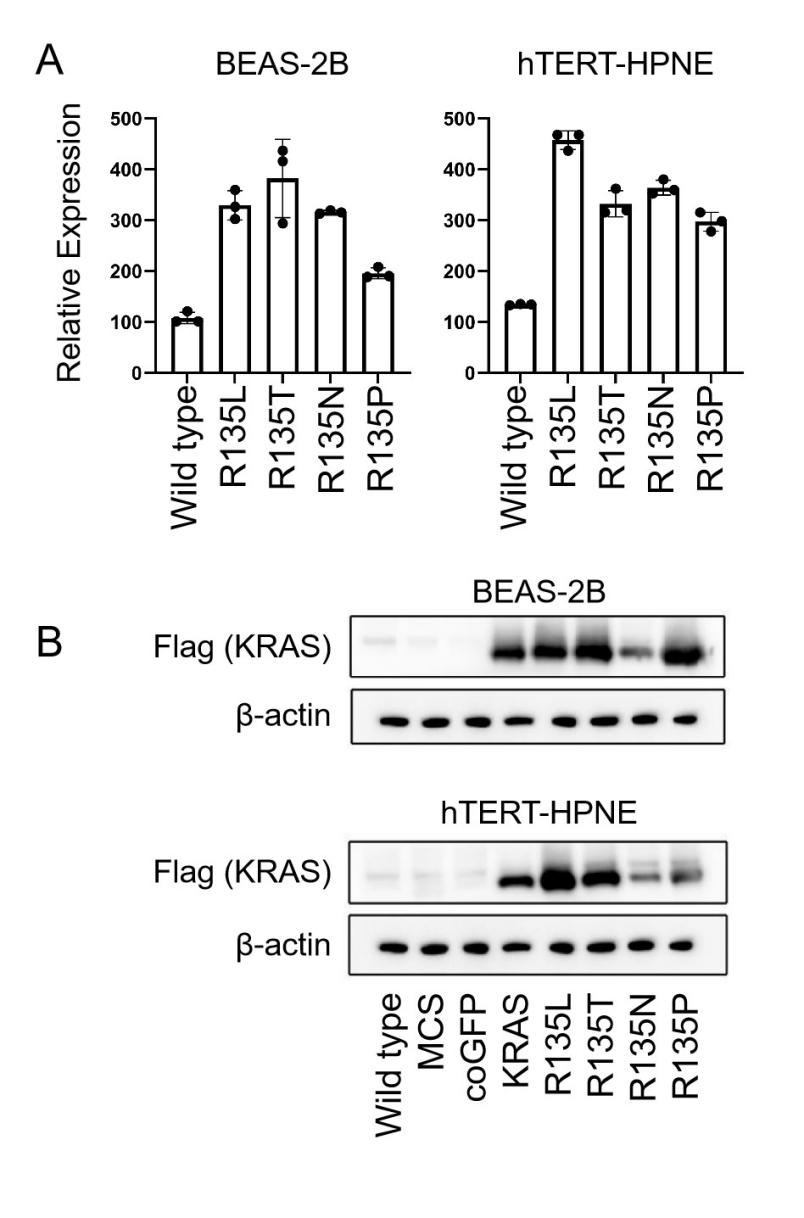

Supplementary Figure 4. Validation of KRAS mutant expression in bronchial and pancreatic epithelial cell lines. A: Quantitative analysis of KRAS transcript levels by RT-PCR. Data are presented as mean±SEM (n=3). B: Western blot analysis confirming the expression of KRAS proteins, with β-actin as control.
